# Genome-wide mapping of EBV-induced genomic variations identifies the role of MUC19 in EBV latency

**DOI:** 10.1101/2024.11.30.625772

**Authors:** Jingwen Yu, Yaohao Wang, Qirong Liu, Xiaohui Zhou, Erle S Robertson, Yonggang Pei

## Abstract

Epstein-Barr virus (EBV) infects over 95% of the world’s population and is tightly associated with multiple human malignant diseases. As the first discovered human oncovirus, EBV is known to induce genomic instability by promoting various types of genomic modifications in host chromosomes. However, the mechanisms through which EBV interacts with the host genome and regulates cellular gene expression in genomic modifications are not yet fully elucidated. In this study, we conducted primary EBV infection in B cells and performed the analyses of copy number variants using whole genome sequencing. After comparing to other distinct B cells with varying EBV infection backgrounds, we identified susceptible regions and unveiled critical host factors induced by EBV infection on chromosomes. Combined with the genome-wide transcriptomic analyses, we found that the mucin gene *MUC19* is distinctively activated by EBV infection and MUC19 promotes oncogenesis through the activation of mTOR signaling in lymphoblastoid cell lines (LCLs). Further results indicate that EBV nuclear antigen 1 (EBNA1) binds to multiple regions within *MUC19* and enhances the expression of *MUC19*. Moreover, sequence alignment has shown intriguing motif homology between the basic repeat unit of *MUC19* and EBV genome, suggesting the potential of EBV-specific integration. To conclude, our study maps the genomic perturbations induced by EBV primary infection and offers new insights into the oncogenic roles of MUC19 during EBV latency.

## Introduction

Epstein-Barr virus (EBV), also known as Human Herpesvirus 4 (HHV-4), infects over 95% of the global population, making it one of the exceptionally successful pathogens^1^. EBV infection is closely associated with a variety of human diseases, including Burkitt’s lymphoma (BL), Hodgkin’s lymphoma (HD), infectious mononucleosis (IM), nasopharyngeal carcinoma (NPC), gastric cancer (GC), and multiple sclerosis (MS)^2,3^. Following initial infection, EBV first infects the host’s oral epithelial cells, and the subsequently released progeny virus particles infect the host B cells, establishing a latent infection inside B lymphocytes in the form of a circular extrachromosomal episome^1^.

Genomic instability is a hallmark of cancer and is characterized by genomic mutations^4^. These aberrant mutations have the potential to trigger tumorigenesis^5,6^. EBV is reported to promote genomic instability in B-lymphocytes^7^. The advancement of high-throughput sequencing methods facilitates the identification of distinct chromosome modifications with higher precision^8,9^. Compared to small fragment genomic variations, structural variations (SVs) bear the potential to further elucidate the chromosomal variations induced by viral antigens. Furthermore, copy number variants (CNVs) typically refer to genomic fragment variations over 1kb in the human genome and are known to contribute to many complex diseases, including neuropsychiatric disorders^10–12^. For instance, a decrease in the copy number of the chemokine gene CCL3L1 significantly increases susceptibility to HIV^13^. Available CNV analyses on the diffuse large B cell lymphoma (DLBCL) and Non-Hodgkin’s Lymphoma (NHL) identified susceptible regions for EBV infection with limited experimental validation^10,14^. Therefore, studying the phenotypic impacts of CNVs in EBV-positive cells has the potential to reveal novel variations and deepen our understanding of EBV-associated oncogenesis.

Regarding EBV infection, previous studies in at least 12 lymphoblastoid cell lines (LCLs) showed that EBV infection induces chromosome modifications across more than 70 chromosomal bands (p < 0.05) with shared integration at 1p31, 1q43, 2p22, 3q28, 4q13, 5p14, 5q12, and 11p15^15^. In Burkitt’s Lymphoma, EBV integration leads to large genomic deletions in the viral genome, including regions of the LMP and EBER genes. Meanwhile, the integrated viral genome disrupts chromosome stability inside infected cells, leading to translocations and deletions on chromosomes 11 and 19^16^. Spectral karyotyping identified frequent structural variations on the host genome that occurred four weeks after post-infection. These mutations stabilized gradually around 25 weeks post-infection, with the final EBV-positive lymphocyte lines exhibiting widespread chromatin and telomere abnormalities^17^. Some EBV-induced LCLs are found to exhibit the classic t(8;14)(q24;q32) translocation, with similar mutations observed in at least five other chromosomes^18^. Collectively, these studies confirm that EBV is capable of interacting with the host genome in lymphocytes and inducing genomic variations, which facilitate DNA damage or repair and reshape B cell genetic landscape. However, the mechanisms by which EBV induces these mutations and their carcinogenic pathways remain largely unexplored.

With the advancement of Next Generation Sequencing (NGS) technologies, whole exome sequencing (WES) on peripheral blood mononuclear cells (PBMCs) and lymphoblastoid cell lines (LCLs) elucidated potential oncogenic single nucleotide polymorphisms (SNPs), along with insertions and deletions^19^. However, the identified SNPs were not further evaluated due to the limitations of sequencing technologies. Also, WES ignores key regulatory non-coding elements such as EBV-specific enhancers. Moreover, the genome-wide association study (GWAS) on 681 clinical NHL patients and 749 controls identified key recurrent mutations in DLBCL within the LOC283177 locus and confirmed mutations associated with chronic lymphocytic leukemia (CLL) at chromosome 13q14, 11q22-23, 14q32, and 22q11.22^20^. Whole genome sequencing (WGS) analysis of NPC patients revealed the role of TGFBR2 in tumorigenesis, providing potential druggable targets for NPC treatment^21^. WGS provides valuable insights into the mutational landscape for studies, but this research highlights the mutational profile of NPC while overlooking the possibility that these mutations could be induced by EBV. These studies, utilizing deep sequencing technologies, confirmed and characterized EBV-induced genomic modifications.

EBV-induced SVs are observed in chromosomes 6, 9, and 15 in both EBV-infected B cells and murine models. Notably, the viral protein BNRF1 also mediates the formation of SV and it is essential for the replication and maintenance of EBV latent infection^7^. EBV proteins such as EBNA1 and BNRF1 can cause chromosomal structural variations, suggesting that variation analysis on larger chromosomal segments might provide new insight to understanding EBV-mediated oncogenesis. EBV episome attached to the host genome through OriP during latency by forming a replication-dependent cross-structure with host DNA^22^. A recent study confirmed the binding of EBNA1 to host chromosomes causes dose-dependent breakage at 11q23 in EBV-positive B lymphocytes, potentially inducing genomic mutation^23^. Their model shows that EBV episome attaches to the human genome at clusters of 18-bp homologous palindrome repeat sequences, resulting in the aggregation of EBNA1 on host chromosome 11q23 region, disrupting chromosomal stability. Using CRISPR-FISH, they observed EBNA1-mediated chromosomal double-strand breakage, and the uneven distribution of chromosome fragments in the next cell cycle, possibly driving EBV-mediated oncogenesis. This study is highly innovative in identifying EBV-induced modifications on chromosome 11q23, while EBV’s ability to induce mutations across the human genome suggests the involvement of additional mechanisms.

Human mucin gene *MUC19* (chr12: 40,393,394-40,570,832), was first cloned and characterized in 2004^24^. *MUC19* is over 177k bp in length and codes a protein of 8384 amino acids with mucin-like threonine/serine-rich repeats in the middle. However, little is known about this gene other than its genomic sequences, particularly in EBV-induced oncogenesis. Here, we conducted an integrated analysis of EBV-related genomic variations across cell lines with different EBV infection backgrounds and highlighted the genomic CNVs induced by EBV primary infection. Furthermore, we identified and characterized the role of the apomucin MUC19 to promote oncogenesis through mTOR activation. Our study bears to potential to provide a theoretical model and experimental procedures for the development of novel antiviral treatment targeting MUC19.

## Results

### EBV induces specific CNVs in B cells

To explore the EBV-induced genomic changes, we performed EBV primary infection in B cells, then collected these B cells with or without EBV infection for whole genome sequencing. We concentrated on analyzing the EBV-induced specific CNVs using the whole genome sequencing data. Our results revealed EBV-induced CNVs across the host genome (Figure 1a), identifying 53 distinct duplications (dups) and 469 deletions (dels), which highlight the genomic instability driven by EBV (Figure 1d, e). Highly mutated CNV duplications were observed at chr17p11.2 (6 dups), chr10p11.1 (4 dups), and chr22q11.1 (4 dups), while frequent deletions were detected at chr11p15.5 (9 dels), chr21q22.3 (8 dels), and chr3p24.3 (8 dels). Compared to previously discovered CNVs in BL and NPC^16,21^, the CNVs shared by EBV-positive B cells are found to overlap with predicted EBV integration sites^25^ and known oncogenic drivers, suggesting the potential oncogenic role of EBV-induced genomic variations (Figure 1a). To characterize the potential role of these CNVs, we next mapped the shared CNVs among the EBV-infected cells (EBVinf), and two EBV-transformed LCLs (LCL1, LCL2) to the high-throughput chromosome conformation capture (Hi-C) data from EBV-positive GM12878 cells, which indicate the active or inactive status of distinct chromosomal regions. The results demonstrate that genomic deletions occur more frequently in transcriptionally active regions in EBV-positive cell lines, while shared segmental duplications are generally within non-coding and intergenic regions (Figure 1b, c).

The identified CNVs also include known oncogenes, such as *DMBT1* and *FOXO1* (Figure 1f, g). Rearrangements within *DMBT1* are frequently discovered in multiple tumors^26,27^. Mutated *FOXO1* is a known driver for decreased overall survival in DLBCL patients and is associated with failure to achieve event-free survival at 24 months (EFS24) after diagnosis^14,28^. Our primary infection induced copy number duplication within *DMBT1* and deletion in *FOXO1*, suggesting their critical functions in EBV infection. Besides CNV analysis, we also performed SNP analysis on EBV-infected primary B cells, which revealed 4,369,022 EBV-related SNP sites. After normalizing the SNP counts according to the lengths of their CNVs, we further defined the 100 most genomically instable CNV regions, overlapping 233 genes. GO enrichment of these high genes shows distinct virus infection and host defense (Figure S1a). In conclusion, we demonstrated the EBV-induced CNVs in the host chromosomes using WGS analyses from primary EBV infection, which suggests the potentially oncogenic roles of the genes within these CNVs.

**Figure 1.**
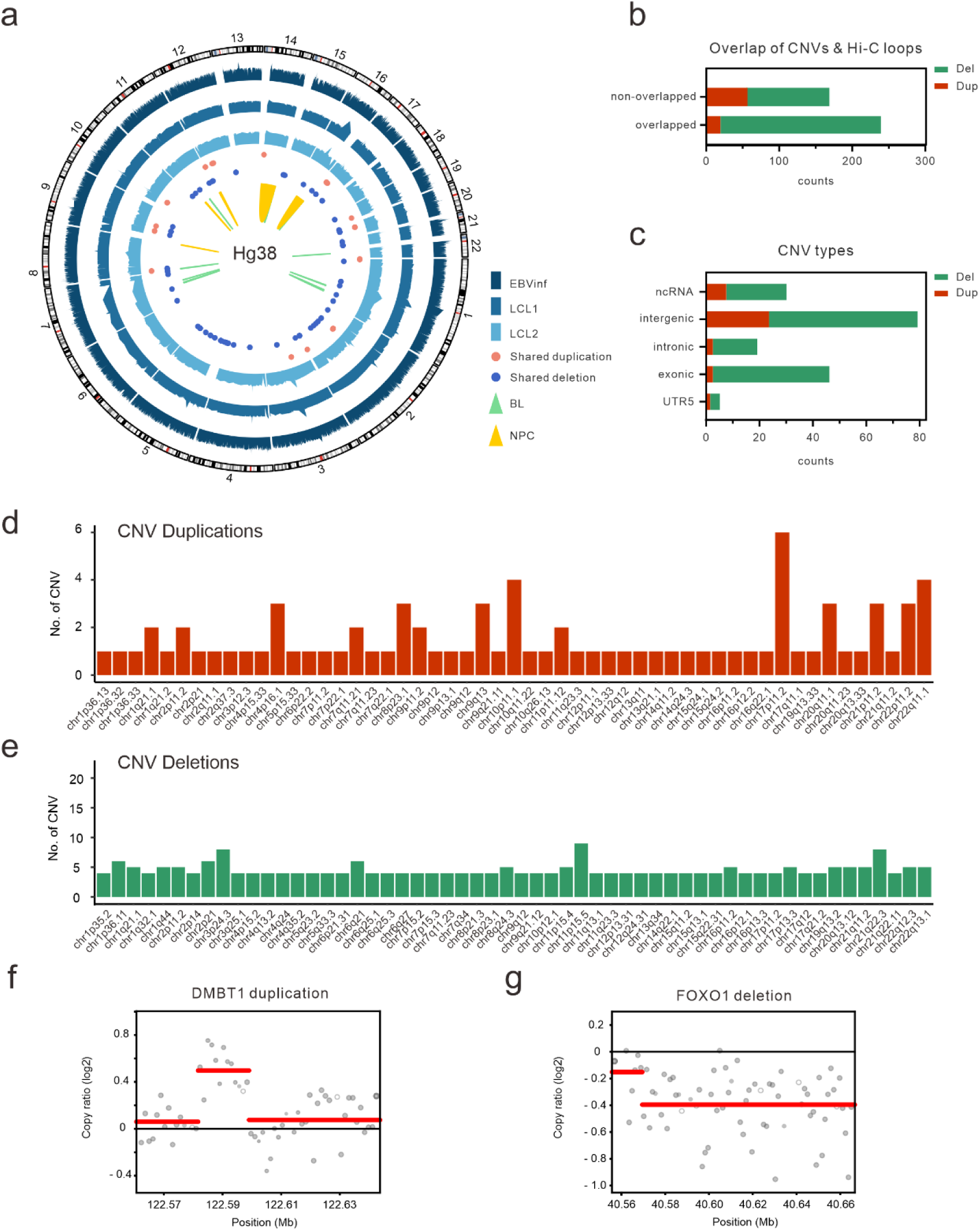
EBV induces specific CNVs in B cells. (a) CNV analysis reveals shared CNVs among EBV-positive cell lines (EBVinf, LCL1 and LCL2). (b) EBV-induced CNVs overlap with actively transcribed genomic regions. (c) EBV-induced CNVs span different genomic regions. (d-e) EBV infection introduces CNV duplications (d) and deletions (e). (f) *DMBT1* is shown to be duplicated as a result of EBV infection. (g) *FOXO1* contains EBV-induced segmental deletion.

### MUC19 shows a high mutation frequency and is distinctively upregulated by EBV infection

To further identify potential key factors associated with the EBV-induced specific CNV regions, we sequenced another two EBV-negative B cell lines (BL41 and BJAB) and then detected the specific CNVs induced by EBV infection across different cellular backgrounds. Comparative analysis between EBV-positive (LCL1 and LCL2) and EBV-negative (BL41 and BJAB) revealed that only one gene, *MUC19*, was susceptible to duplication in EBV-positive B cells, while seven other genes were found to be susceptible to deletion (Figure 2a-c; S2a). The following genomic qPCR results confirmed the indicated CNVs across *MUC19* DNA in several EBV-positive cell lines (Figure 2d, e). Furthermore, real-time qPCR also showed that *MUC19* expression was significantly higher in EBV-positive cell lines (LCL1, LCL2, GM12878) compared to the EBV-negative cell lines (Ramos, BJAB, DG75), which suggests that the expression of MUC19 links to EBV latent infection (Figure 2f).

Duplicated genes within CNV regions are shown to regulate pathogenesis by overexpression through dosage effect^29^. To further explore the expressional changes in CNV-associated genes during EBV primary infection, we analyzed transcriptome sequencing of EBV-infected primary B cells (Figure 2g; S2b)^30^. The results indicated that *MUC19* exhibited a significant upregulation in EBV-infected B cells 14 days post-infection, along with other members of the mucin family, MUC1 and MUC12 (Figure 2g). The following KEGG analysis showed multiple signaling pathways tightly associated with EBV primary infection, including cell cycle, NF-κB signaling, and TNF signaling (Figure 2h).

**Figure 2.**
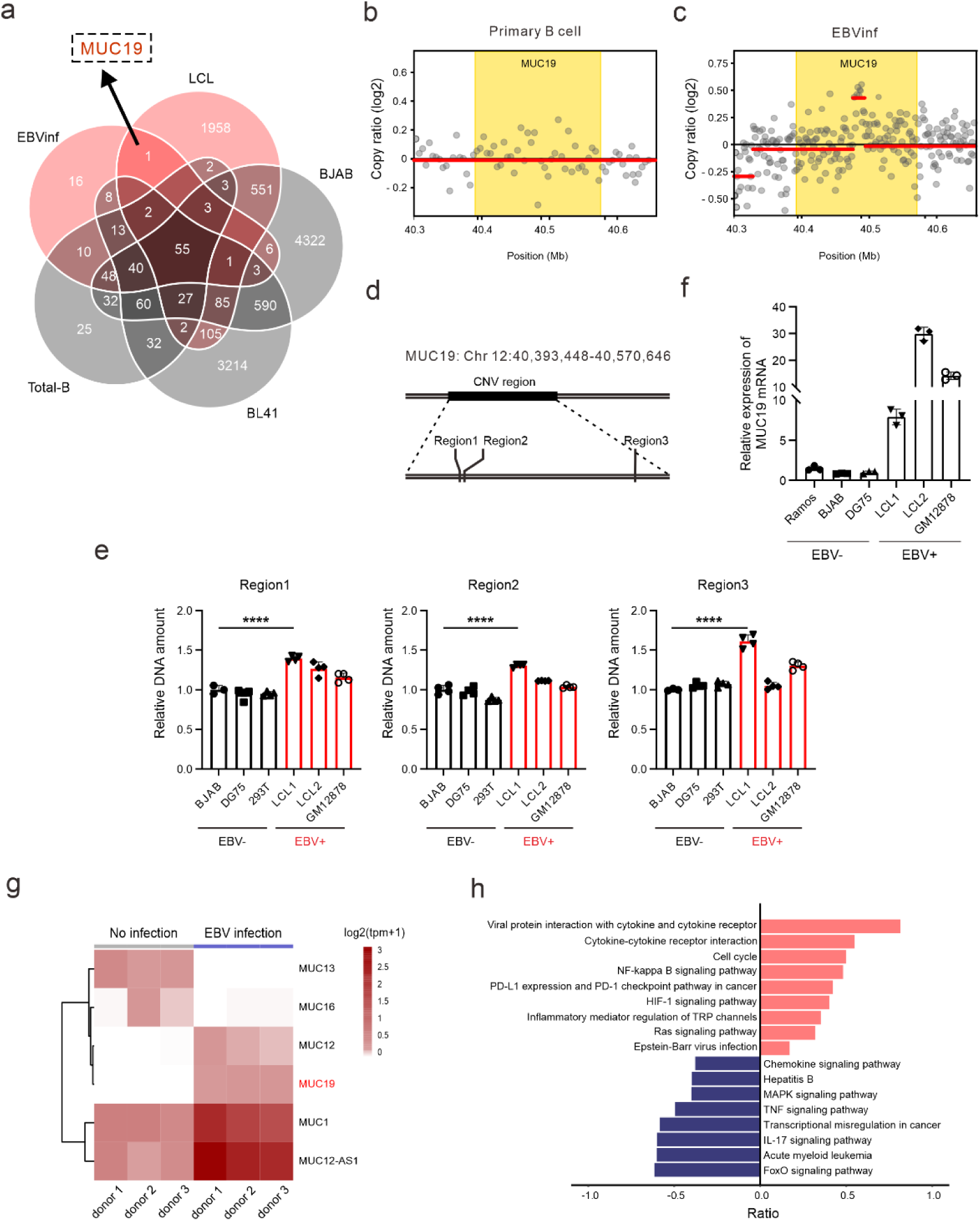
MUC19 shows a high mutation frequency and is distinctively upregulated by EBV infection. (a) The comparative analysis identifies shared CNV duplications within *MUC19* among EBV-positive cell lines. (b-c) *MUC19* is shown to be duplicated by EBV primary infection. (d-e) Genomic qPCR confirms the CNV identified by WGS in EBV-positive cell lines. (f) The qPCR result suggests active MUC19 expression in EBV-positive cell lines. (g) Multiple mucin factors are differentially expressed by EBV infection. (h) KEGG analysis reveals cellular signaling pathways regulated by EBV primary infection.

### MUC19 promotes cell cycle in EBV-positive B cells

As typical secreted proteins, MUC19 has 3 von Willebrand factor D (VWD) domains near its N-terminus, and a von Willebrand factor C (VWC) domain followed by a C-terminal cystine-knot (CTCK) domain^31^ (Figure 3a). Previous studies showed *MUC19* secretion in glandular tissues and epithelial cells including major salivary glands^24^, lacrimal glands^32^, middle ear epithelium^33^, and airway tissues^34^. However, our results indicate that MUC19 localizes in the cytoplasm with no observable secretion for lymphoma cell lines (Figure 3b; S3b). MUC19 shares close homology with MUC2, MUC6, MUC7, MUC5A and MUC5B (Figure S3a). Proteomic analyses indicate spontaneous cleavages within MUC5AC and MUC5B^35–37^, and the cleavages are also identified in MUC2 and MUC3 in epithelial cells^38,39^. We discovered a 25kD cleavage in EBV-positive LCLs using a C-terminus-derived antibody, which appears to be in proportion to the overall mRNA amount (Figure 2f; S3c).

Next, we explore the oncogenic role of MUC19 in B lymphoma cells. Cell proliferation assays showed that lymphoma MUC19 knockdown cell lines demonstrated significantly reduced cell viability (Figure 3c-f). The abrogated viability suggests the importance of MUC19 in promoting cell cycle and EBV-induced oncogenesis. MUC19 knockdown in LCL1 exhibits cell cycle arrest before the replication and triggers apoptosis, further supporting its oncogenic role in EBV latency (Figure 3g, h; S3d). Moreover, our study reveals a decreased expression of Cyclin D1 in MUC19 knockdown LCL1 cells, indicating the potential role of MUC19 in facilitating the cell cycle by regulating Cyclin D1 (Figure 3e).

**Figure 3.**
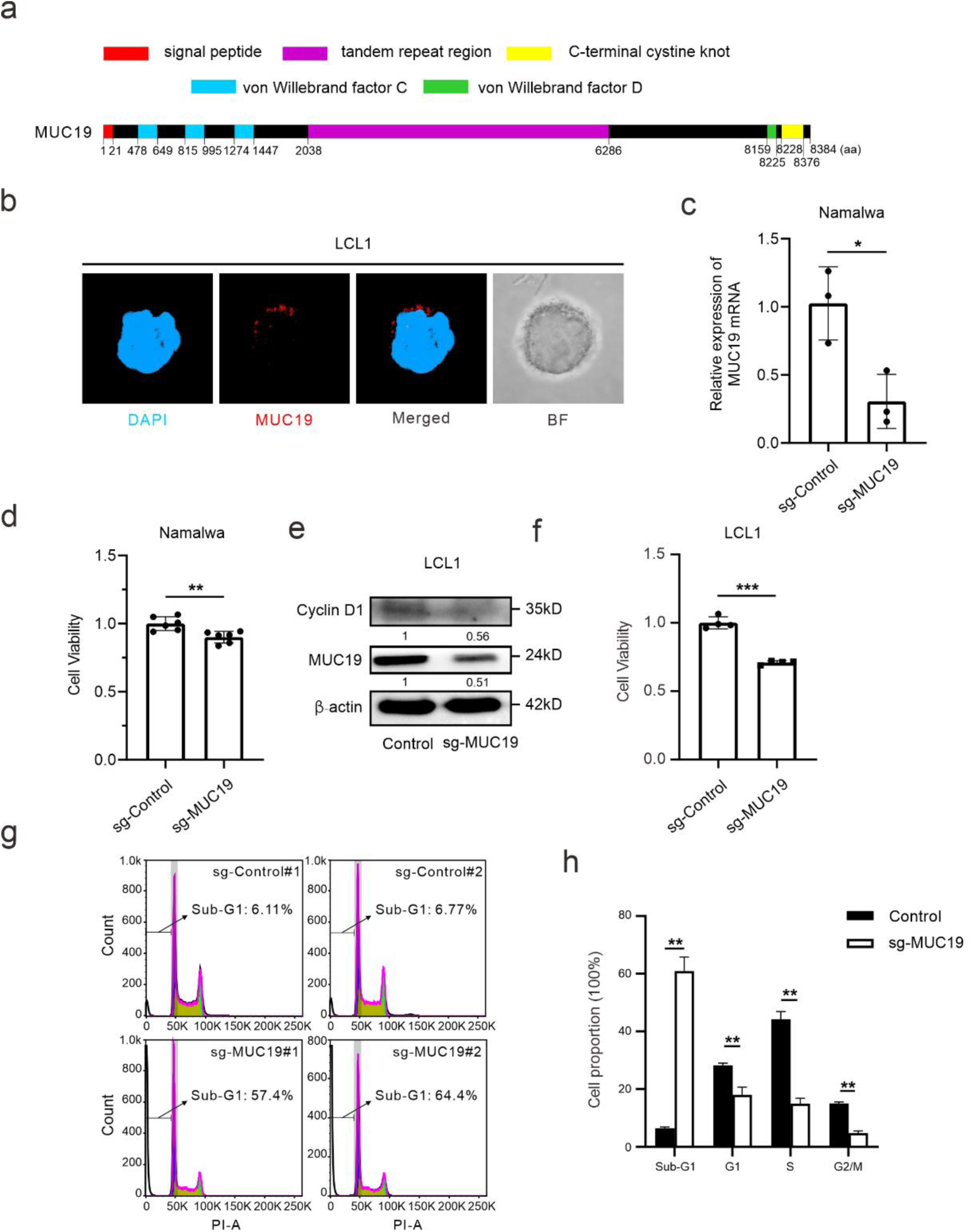
MUC19 promotes cell cycle in EBV-positive B cells. (a) A schematic diagram illustrates the structure of MUC19 protein. (b) Immunofluorescence imaging indicates that MUC19 localizes in the cytoplasm in B cells. BF, bright field. (c-d) MUC19-knockdown cells exhibit abrogated viability in Namalwa. (e-f) MUC19-knockdown cells exhibit abrogated viability that potentially resulted from the down-regulation of Cyclin D1. (g-h) MUC19-knockdown LCL1 shows intense apoptosis. **, p < 0.001.

### MUC19 promotes B cell proliferation by activating the mTOR pathway

Previous CNV analysis identifies MUC19 as a highly mutated and promising novel diagnostic marker for Hepatoid Adenocarcinoma of the Stomach (HAS)^40^. Moreover, overexpression of MUC19 points to nuclear localization of β-catenin and Wnt antagonist abrogated the expression of downstream Wnt pathway markers Cycin D1 and c-Myc. However, direct evidence underlying how MUC19 promotes the oncogenic pathways is still lacking. The size and structure of MUC19 (8384aa) present great difficulty in studying its function via traditional molecule techniques (Figure 3a). Thus, we first explored its function by focusing on its specific domains. A study indicates that the C-terminus of MUC1 and β-catenin increased SNAIL transcriptional activity to activate epithelial-mesenchymal transition (EMT) in cancer cells^41^. However, we did not observe the significant effects in Wnt/β-catenin, EMT, and the MAPK cascade when the VWD and CTCK domains of MUC19 protein were overexpressed.

To further explore the functions of MUC19 in EBV latency, we focused on the repeated region of MUC19, which contains approximate tandem repeats of “GVTGTTGPSA” (Figure 3a). The motif analysis revealed approximately 420 copies of this basic repeat unit on *MUC19* (5’-GGAGTGACAGGGACAACTGGACCATCAGCT-3’, p < 0.00002) (Figure 4a). To validate the biological function of the tandem repeats, we overexpressed the seemingly conserved basic unit in a single copy (Figure 4c, d). The results showed that one copy of the basic repeat unit promotes cell viability and triggers the phosphorylation of phosphoinositide 3-kinase (PI3K) and mTOR in 293T and Akata cells, activating the transcription of downstream factors such as Cyclin D1 (Figure 4e-h). These results suggest the oncogenic role of MUC19 in activating mTOR signaling to promote EBV-mediated latency and oncogenesis.

**Figure 4.**
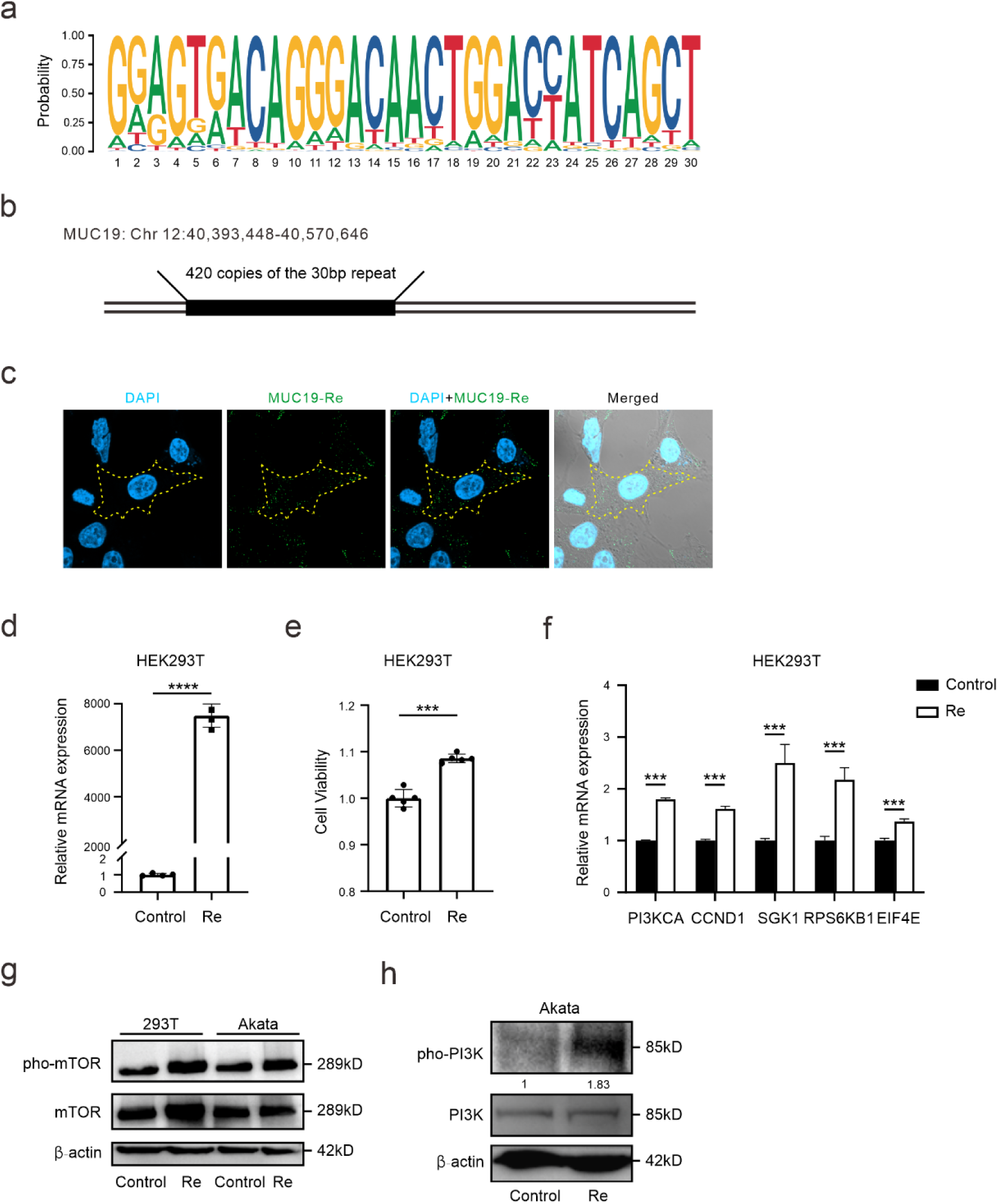
MUC19 promotes B cell proliferation by activating the mTOR pathway. (a) Motif analysis revealed that a conserved 30-bp sequence is the basic unit that encodes the repeat region of MUC19. (b) *MUC19* is shown to contain 420 copies of the 30-bp repeat (p < 0.00002). (c-d) The basic repeat unit is successfully transfected into HEK293T cells. (e) The basic repeat unit is shown to promote cell proliferation and growth. (f) RT-qPCR indicates the basic repeat unit promotes mTOR activation. (g-h) The basic repeat unit is indicated to increase the phosphorylation of mTOR and PI3K. Re, the 30-bp basic repeat unit.

### EBNA1 binds to MUC19 gene and enhances its expression

EBV nuclear antigens are shown to disrupt host genomic stability, potentially leading to oncogenic mutations^42^. Among them, EBNA1 is a multi-functional EBV-encoded latent protein that serves as a transcription activator^43^. EBNA2 and EBNALP are also EBV-encoded transcription factors^44^. Expression of these antigens revealed that EBNA1 significantly upregulates the expression of MUC19 in 24 hours post-transfection (Figure 5a, b). EBNA1 also activates Cyclin D1 expression, suggesting that this activation may be mediated through MUC19 (Figure 5c).

EBNA1 is shown to aggregate around certain chromosomal regions and triggers double-strand breakage^23^. Next, to explore whether EBNA1 regulates MUC19 expression through binding, we performed the ChIP assays in two EBV-positive cell lines (Akata and LCL1) and confirmed that EBNA1 can bind to the MUC19 promoter and its multiple genomic regions, potentially upregulating MUC19 expression as a transcription factor (Figure 5d, e). Additionally, the ChIP assay reveals the active binding of EBNA1 to the MUC19 repeat region, suggesting the tight interaction between EBNA1 and multiple repeat regions of *MUC19* (Figure 5f). Together, these experiments highlight the strong association of EBV-encoded EBNA1 and MUC19, emphasizing the critical role of MUC19 in EBV-mediated oncogenesis.

Furthermore, multiple sequence alignment indicates homology between the repeat regions and the EBV genome (Figure 5g-j). The homologous sequence most specifically entails 19bp of the total 30bp sequence and points to the intrinsic repeat in EBV genome, such as BNRF1, EBNAs, BHLF1, BMRF1, LF3, and so on (Figure 5j). This interesting homology suggests a potential recombination of EBV and the host genome, specifically MUC19, during EBV evolution. In the EBV genome, 51 homologous occurrences were discovered (p < 0.00005). In the MUC19 DNA sequence, 351 motif occurrences were discovered (p < 0.00005), while in the entire human genome (hg38), 213 strict matches were discovered (p < 1e-8). 86 of those are within MUC19 (chr12:40,393,394-40,570,832) and 8 are located in the gene TBCD (chr17:82,752,042-82,945,914). The rest are distributed randomly across the whole genome.

To conclude, we have explored the roles of MUC19 in promoting cell cycle and oncogenesis, focusing on its repeat region to activate the mTOR pathway. Combined with our primary infection and WGS analysis, we demonstrate that EBV infection promotes genomic instability with EBNA1, which triggers CNV at the MUC19 locus and activates its expression. The expressed MUC19 then activates mTOR signaling, promoting cell proliferation, growth, and survival (Figure 5k).

**Figure 5.**
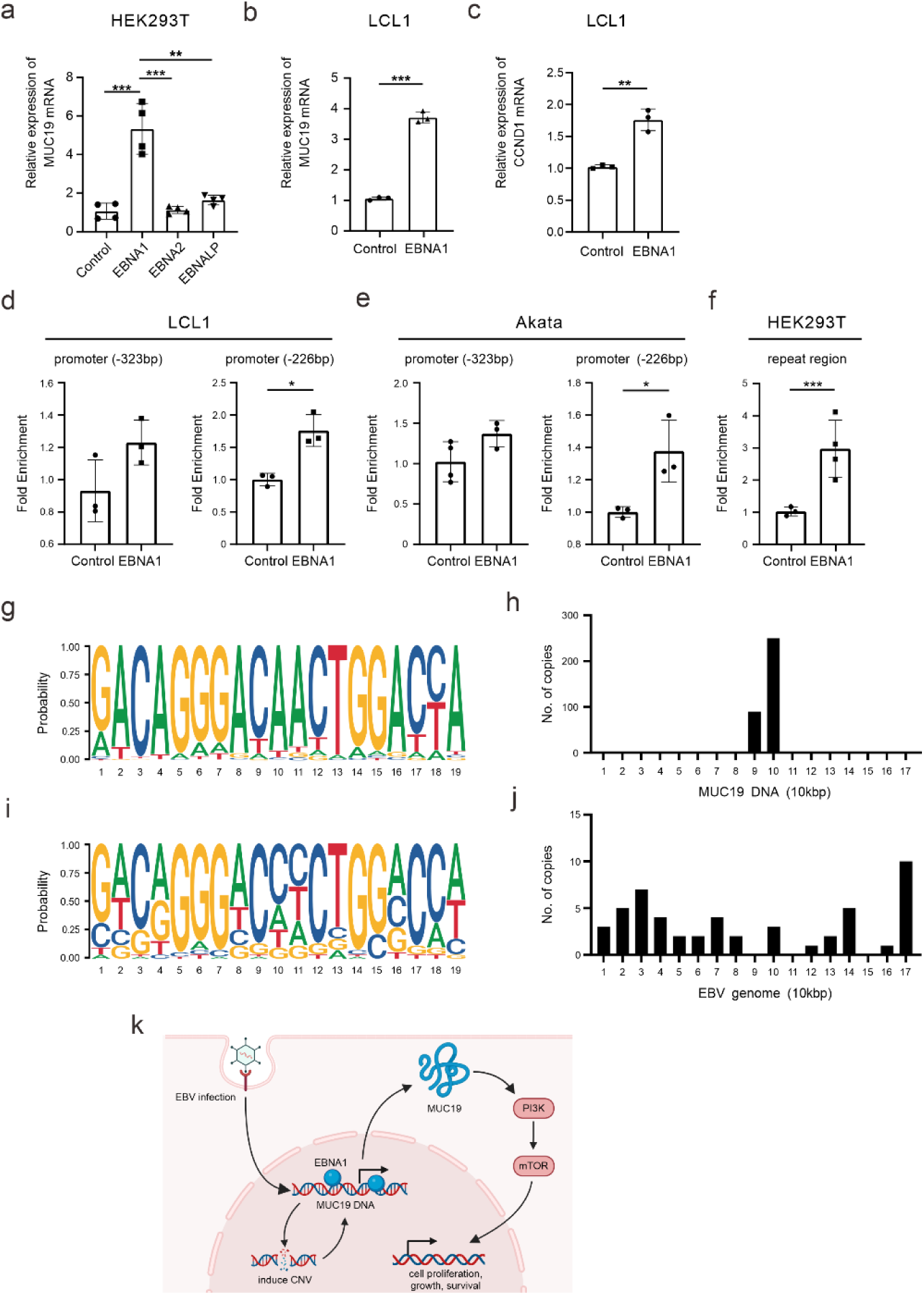
EBNA1 binds to MUC19 gene and enhances its expression. (a-b) EBNA1 activates the expression of MUC19 in HEK293T (a) and LCL1 (b). (c) EBNA1 promotes CCND1 expression in LCL1. (d-e) ChIP assays show that EBNA1 binds to the promoter region of *MUC19* in LCL1 (d) and Akata (e). (f) ChIP assay indicates that EBNA1 actively binds with the repeat region of MUC19. (g) Sequence alignment discovered a 19-bp motif in the MUC19 repeated region. (h) The motifs are enriched in the CNV region of MUC19. (i) The corresponding motif is found in the EBV genome. (j) The locations of the motifs are dispersed throughout the EBV genome. (k) The graphical abstract shows the model of this study, which was created with BioRender (biorender.com).

## Discussion

Our study found the shared copy number duplication of MUC19 induced by EBV primary infection and in EBV-positive cell lines. We then validated the high MUC19 expression in EBV-positive LCLs, suggesting the functional roles of MUC19 in EBV oncogenesis. We next identified the functional regions of MUC19 to promote oncogenesis via activating the mTOR pathway. Finally, we confirmed the relationship between EBNA1 and MUC19.

Although CNV is pervasive in EBV-positive cell lines, the expressional changes brought about by the CNV dosage effect do not fully account for the expressional changes. The up-regulation of MUC19 is observed in HEK293T cells in only 24 hours post EBNA1 overexpression, while upregulation induced by the CNV dosage effect requires a longer time. Therefore, these results suggest that EBNA1 serves as a transcription activator to promote MUC19 expression. Besides MUC19, there were other interesting shared CNVs from our primary infection, including IRF4/DUSP22 duplication and NOTCH2NLA deletion (Table S1). Interestingly, IRF4 and DUSP22 are within the same EBV super enhancer^45^. qRT-PCR result indicates DUSP22 activation in EBV-positive B-cells, suggesting a potential functional role in EBV latency (Figure S2c). NOTCH2NLA (Notch Homolog 2 N-Terminal-Like Protein A) is downregulated in LCLs, possibly due to EBV infection-induced copy number deletion (Figure S2d). However, NOTCH2NLA activates the NOTCH pathway and facilitates EBV oncogenesis. The expressional change brought by CNV cannot provide a functional explanation for EBV-induced NOTCH2NLA deletion. Further experiments are required to further explore the role of NOTCH2NLA in EBV-related malignancies.

The MUC19 region is also associated with multiple EBV-related transcription factors. Chromatin immunoprecipitation sequencing (ChIP-seq) analyses for several EBV-associated transcription factors in the EBV-positive GM12878 cells show active binding at the MUC19/LRRK2 locus, such as CTCF and POLR2A, indicating active transcription (Figure S2e). H3K4me1 signal shows that the MUC19/LRRK2 gene and the adjacent CNTN1 are located within a closely connected topological domain, suggesting co-regulation. Other transcription factors have also been found to bind to this gene locus, including EBV-associated IRF4, PAX5, BCL3, RUNX3, and NFAT. These results suggest that MUC19 is tightly involved in EBV-mediated latency and oncogenesis.

Although MUC19 is the only mucin factor identified from our screen with copy number duplication, mucin factors in general are differentially regulated upon EBV infection (Figure 2g). For instance, the membrane-tethered MUC12 exhibited a similar expression pattern as MUC19 after EBV primary infection. The oncogenic role of MUC12 in renal cell carcinoma is related to the c-Jun/TGF-β signaling pathway^46^. As the mucin family is tightly associated with numerous inflammatory pathways and immune functions of various pathogenesis, these genes could also participate in the EBV-associated biological processes. Future efforts may explore the roles of other mucin factors in regulating EBV-mediated diseases.

The mTOR pathway is essential for cell growth and human tumorigenesis^47^. Previous study suggests mTOR activation is mediated in the transformed B cells independent of the PI3K/Akt signaling pathway^48^. In our functional experiments, we have confirmed the mTOR activation through a pathway independent of Akt, and another PI3K pathway in which SGK1 compensates for Akt and activates the downstream factors including S6K1, EIF4E, and CCND1 (Figure 4f).

In conclusion, our study combined WGS data from EBV-transformed primary B cells and other LCLs to characterize EBV-induced chromosomal copy number variations. Integrative analysis and expressional assays identified MUC19 as a critical factor for EBV infection. Despite the hardship of studying mucin factors, we managed to unveil the role of MUC19 in regulating cell cycle to promote oncogenesis through mTOR activation. Furthermore, we revealed the functional domain of MUC19 and showed its close interaction with EBV-encoded EBNA1 antigen. Our study further elucidated the genomic mutations induced by EBV and highlights the potential of MUC19 as a novel therapeutic target for anti-EBV treatment.

## Materials and Methods

### Ethical statement

The naïve B cells from healthy donors in this study were provided by the University of Pennsylvania Immunology Core (HIC) with written and informed consent. All the procedures were approved by the Institutional Review Board (IRB) and conducted according to the declarations of Helsinki protocols^49^.

### Plasmids

pMD2.G (Addgene #12259) and PsPAX2 (Addgene #12260) are used for lentivirus packaging. EBV latent genes are cloned into the pEGFP-C1 (Takara, Japan). The Re plasmid in the study is generated by cloning the basic repeat unit of MUC19 into the pCMV-Myc-N vector (Takara, Japan). For CRISPR systems, pLG15-NC containing control sgRNA is a generous gift from Dr. Ruilin Tian (Southern University of Science and Technology, China). The plasmid dCas9-KRAB-MeCP2-GFP is modified from lenti_dCas9-KRAB-MeCP2 (Addgene #122205) by replacing the BSD with GFP, which is also from Dr. Ruilin Tian. The sgRNA primers used for CRISPR inhibition and knockout systems are designed using CHOPCHOP (chopchop.cbu.uib.no).

### Cell culture

HEK293T (human embryonic kidney cell line) cells were cultured in Dulbecco’s modified Eagle’s medium (DMEM) medium supplemented with 10% fetal bovine serum, 50 U/ml penicillin, 50 μg/ml streptomycin, and 2 mM L-glutamine. EBV-negative cells (BL41, BJAB, and Ramos) and EBV-positive cells (Namalwa, Raji, LCL1, LCL2, and GM12878) were cultured in RPMI 1640 medium supplemented with 10% fetal bovine serum, 50 U/ml penicillin, 50 μg/ml streptomycin, and 2 mM L-glutamine. All the cells were cultured in a 37 ℃incubator at 5% CO^2^.

Transfections into HEK293T were performed using a Calcium Phosphate Cell Transfection Kit (C0508, Beyotime, China). Transfections into LCL1 and Akata are conducted using Gene Pulser Xcell Electroporation Systems (Bio-Rad, USA). CRISPR knockout in LCL1 is performed using lentiCRISPRv2 (Addgene #98290) and selected with puromycin at 0.5 μg/ml for 3 days. Surviving cells were then grown in a normal medium for amplification.

### EBV primary infection and whole genome sequencing

Approximately 10 million B cells were infected with EBV for 15 days, while 5 million B cells were stored at -140 degrees as negative control. Then both of the infected or uninfected B cells were collected, and the genomic DNA was extracted from these obtained cells to establish a library using Nextera DNA Flex Library Preparation kits (Illumina, USA). Subsequently, the libraries were submitted to the Genome Technology Access Center (GTAC) at Washington University in St. Louis for whole genome sequencing using the NovaSeq 6000 system. Besides the collecting cells in EBV primary infection, other EBV-negative or EBV-positive cells were also sent for whole genome sequencing with the same procedures.

### Multi-omics Sequencing Data Analysis

CNVkit (0.9.4)^50^ was used to call CNVs from WGS data. Primary B cell WGS data was used as normal tissue reference for the EBV-transformed cell line CNV analysis. CNVs of other cell lines are called against a flat reference. Comparative analysis was performed using bedtools (2.30.0) and R scripts. Visualizations are made using the R package circlize (0.4.16). The WGS data underwent standard processes for quality control and alignment to human genome assembly Hg38.

Somatic SNVs and small insertions and deletions (indels) were performed using GATK^51^. Joint analysis of CNV and SNV is done by intersecting SNV loci with CNV segments using the R package GenomicRanges (1.52.0). The CNV regions were sorted in descending order based on the number of SNVs contained within each region. To normalize the data, mutation scores were calculated by dividing the number of SNVs, and normalized by the length of the corresponding CNV region. The top 100 CNV segments with the highest mutation scores were selected for enrichment analysis.

RNA-seq data was analyzed using patient data from primary B cells infected with the Epstein-Barr virus on Day 0 and Day 14. Raw data is available at the Gene Expression Omnibus (GEO) under accession number GSE125974^30^. The raw data of gene expression were quantified using Salmon (0.8.2). Differentially expressed genes were obtained using the DESeq2 (1.40.2) in R, with p-value < 0.05 and absolute fold change > 5. KEGG analysis is carried out using the R package clusterProfiler (4.8.3), and visualized by ggplot2 (3.5.1). ChIP-seq and Hi-C data were obtained online from the EBV-positive GM12878 through the ENCODE database (encodeproject.org).

### Quantitative real-time PCR

Quantitative real-time polymerase chain reaction (qRT-PCR) was utilized to measure the expressions of the target genes. Total RNA was extracted using an RNA extraction kit (Beyotime, China) following the manufacturer’s protocol. The quality and quantity of the RNA were assessed using a spectrophotometer, and only samples with high-quality RNA (A260/A280 ratio between 1.8 and 2.0) were used for cDNA synthesis. The synthesized cDNA was then used as a template for qRT-PCR amplification using gene-specific primers and a SYBR Green master mix (Yeason, China). The qRT-PCR reactions were run on a real-time PCR machine with the following cycling conditions: initial denaturation at 95°C for 5 minutes, followed by 40 cycles of denaturation at 95°C for 15 seconds, annealing at 55°C for 30 seconds, and extension at 72°C for 30 seconds. The relative expression levels of the target genes were calculated using the 2^(-ΔΔCt) method, normalized to an internal control gene GAPDH. All qRT-PCR experiments were performed in at least triplicate to ensure reproducibility and statistical analysis was conducted to determine the significance of the results.

### Cell Viability Assay

Cell viability was measured using the Cell Counting Kit-8 (Beyotime, China). Cells were transferred to a 96-well plate at a density of 5,000 cells per well in 100 µL medium. 10 µL of CCK-8 solution was then added to each well. Then the optical density (OD) at 450 nm was measured 2 hours after the addition of the CCK-8 reagent using BioTek Synergy H1 microplate reader (Agilent Technologies, USA).

### Immunofluorescence

Adherent cells were grown on coverslips mounted on a 6-well plate before CRISPRa transfection at the latest passage. Suspension cells are centrifuged after 48h CRISPRa plasmid transfection and then fixed on coverslips with 4% paraformaldehyde, permeabilized with 0.1% Triton X-100 for 10 minutes, and blocked with 5% bovine serum albumin (BSA) for 20 minutes at room temperature. Primary antibodies against the target protein were then applied and incubated overnight at 4°C. After washing with phosphate-buffered saline (PBS), cells were incubated with fluorochrome-conjugated secondary antibodies for 1 hour at room temperature in the dark. Nuclei were counterstained with DAPI. Fluorescent images were taken using a confocal microscope LSM 900 (Zeiss, Germany) equipped. Image processing was performed using Zen blue (3.4.91).

### Western Blot and ELISA

WB experiments are a technique commonly used to detect protein levels. First, proteins are extracted using cell lysis buffer, and after measuring protein concentration, proteins are separated by electrophoresis on an SDS-PAGE gel. Then, proteins are transferred to a PVDF membrane, and non-specific binding sites are blocked with 5% skim milk, followed by overnight incubation with a specific primary antibody. After washing the membrane, it is incubated with HRP-labeled secondary antibody for 1 hour, and the membrane is washed again. Finally, protein signals are detected through chemiluminescence, and the intensity of protein bands is quantitatively measured using the image analysis software ImageJ (1.54). ELISA is performed to evaluate the secretion of MUC19 in B lymphoma cells using a commercially available ELISA kit (CUSABIO, China).

### Flow Cytometry

Apoptosis was detected using the Cell Cycle and Apoptosis Analysis Kit (Beyotime, China), which employs the propidium iodide (PI) staining method for cell cycle and apoptosis analysis. After staining the cellular DNA with propidium iodide, flow cytometry FACSCanto SORP (BD Biosciences, USA) is used to analyze cell cycle and apoptosis using FlowJo (10.6.2).

### Motif Discovery for the consensus repeat

The motif discovery was conducted using FIMO (5.5.5)^52^. The p-value is defined as the probability of a random match for the desired sequence with as good or better a score. The score is computed by summing the appropriate entries from each column of the position-dependent scoring matrix that represents the motif. R package ggseqlogo (0.2) was used to visualize the motifs.

## Supplementary Materials

**Figure S1.**
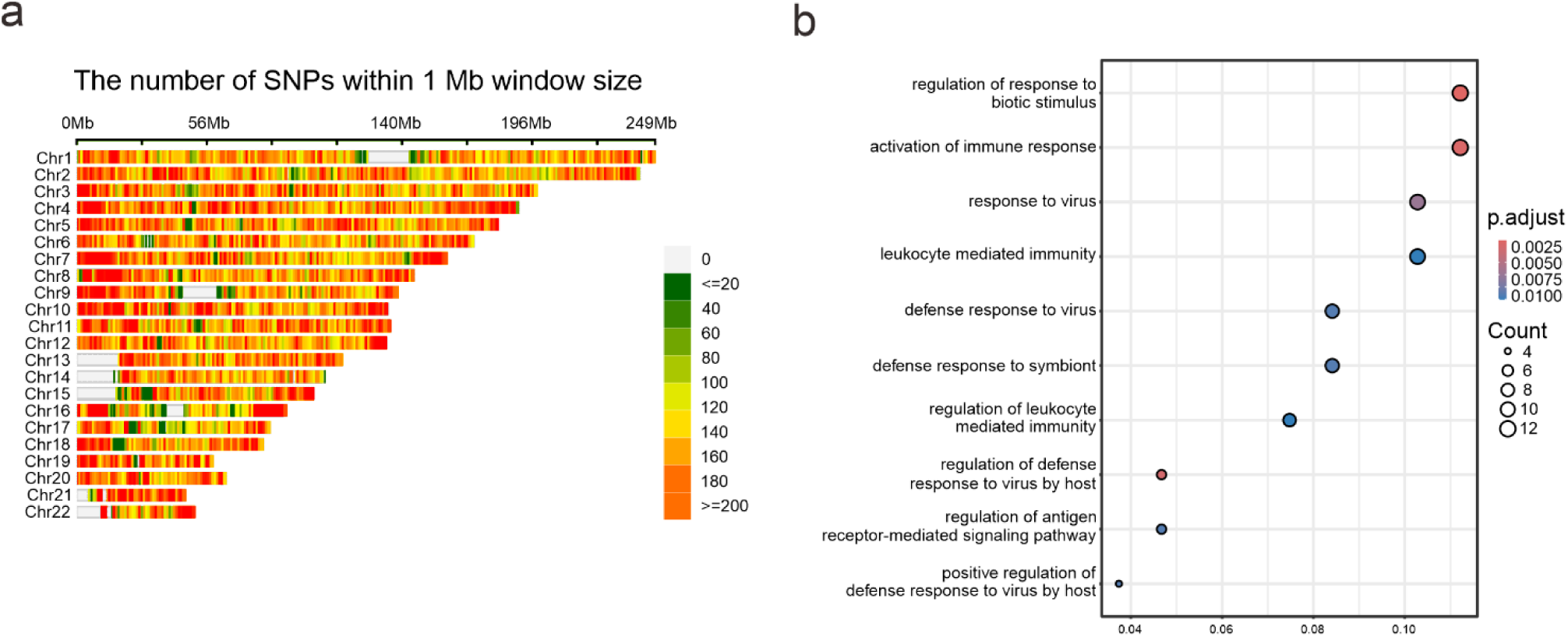
(a) EBV introduces SNPs across the human genome. (b) The highly mutated regions are shown to possess anti-viral factors.

**Figure S2.**
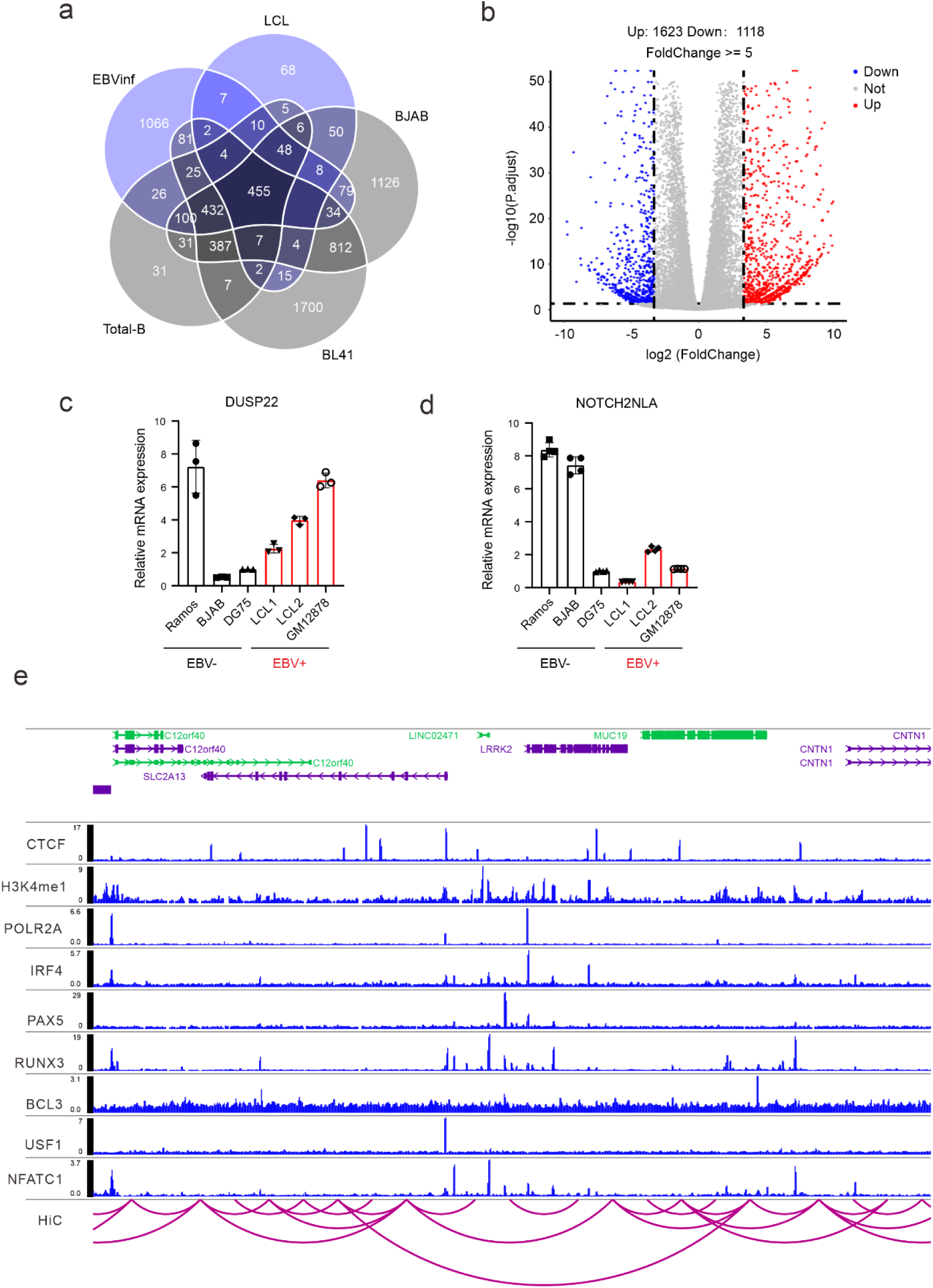
(a) Comparative analysis reveals shared genomic CNVs shared among EBV-positive cell lines. (b) RNA-seq analysis shows differentially expressed genes by EBV primary infection. (c) DUSP22 is differentially expressed in B cells. (d) NOTCH2NLA is silenced by EBV infection. (e) The LRRK2/MUC19 region is highly regulated by EBV-related transcription factors.

**Figure S3.**
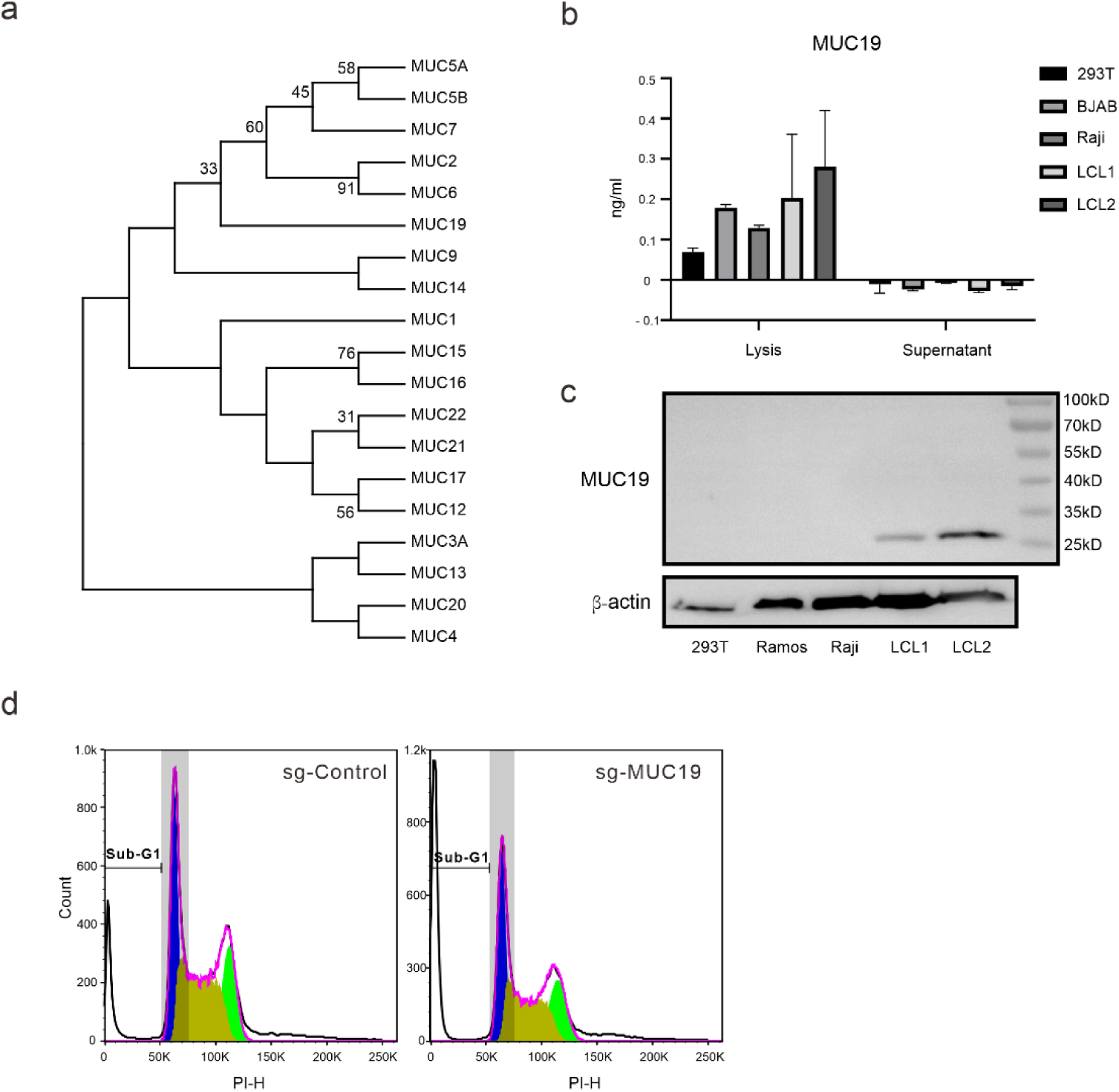
(a) Comparative genomics shows the homology among all identified mucin factors. (b) ELISA assay suggests that MUC19 is not secreted in B cells. (c) Western blotting identifies a 25-kD C-terminal cleavage in EBV-positive cells. (d) Apoptosis assay replicated in Namalwa cell lines indicates that MUC19 promotes cell proliferation and survival.

## Acknowledgment

We are grateful to Dr. Zhi Wei (New Jersey Institute of Technology) and Dr. Tian Tian (Wuhan University) for their technical support and valuable suggestions.

